# The hypothalamic paraventricular nucleus mediates vagus nerve stimulation-enhanced gastric emptying

**DOI:** 10.64898/2026.10.22.758166

**Authors:** Chun-Huei Zou, Yu-Chieh Huang, Wu-Lung Tseng, Ye-Ting Lin, Ying-Jui Chao, Yi-Jen Wu

**Affiliations:** Institute of Clinical Medicine, College of Medicine, National Cheng Kung University, Tainan 70101, Taiwan; Division of General Surgery, Department of Surgery, National Cheng Kung University Hospital, College of Medicine, National Cheng Kung University, Tainan 70403, Taiwan; Department of Neurology, National Cheng Kung University Hospital, College of Medicine, National Cheng Kung University, Tainan 70101, Taiwan

**Keywords:** Vagus nerve stimulation, Hypothalamic paraventricular nucleus, Gastroparesis, Gastric emptying, Motilin, Anti-inflammatory effects, Brain–vagal–gut axis

## Abstract

**Background:** Vagus nerve stimulation (VNS) improves gastrointestinal (GI) motility, but its central regulatory mechanisms remain unclear. The hypothalamic paraventricular nucleus (PVN) regulates autonomic and neuroendocrine functions. We investigated whether PVN centrally regulates the prokinetic effects of VNS.

**Methods:** Cervical VNS was applied to rats with intestinal manipulation (IM)-induced gastroparesis. GI motility was assessed by gastric emptying (GE) and geometric center (GC) of fluorescein isothiocyanate–dextran. PVN activation during VNS was evaluated by extracellular neuronal firing recording and c-Fos immunostaining. PVN electrical stimulation and muscimol inhibition assessed its causal role. Serum levels of GI motility-related hormones ghrelin and motilin, inflammatory markers TNF-α, interleukin (IL)-6, and corticosterone, and gut expression of motilin receptor (MTLR) and myeloperoxidase (MPO), a neutrophil marker, were assessed as potential downstream mediators.

**Results:** IM impaired GE and GC and increased TNF-α and IL-6. Cervical VNS enhanced GE and GC, reduced TNF-α, IL-6, and MPO, and activated the dorsal motor nucleus of the vagus, nucleus of tractus solitarius, and PVN. PVN neuronal firing increased during VNS in a stimulation-strength dependent manner. Direct PVN stimulation at VNS-evoked firing frequencies enhanced GE, MTLR expression, and serum motilin, which positively correlated with GE. PVN stimulation also reduced IL-6 and MPO. Conversely, PVN inhibition attenuated VNS-enhanced GI motility, and MTLR expression and reduced its anti-inflammatory effects on IL-6 and MPO, although IL-6 was not correlated with GE.

**Conclusion:** PVN acts as a central regulator within the vagal circuit, activated by VNS and gating VNS-enhanced vagal efferent output for gastric emptying and anti-inflammatory effects.

PVN mediates VNS-enhanced gastric emptying in experimental gastroparesis

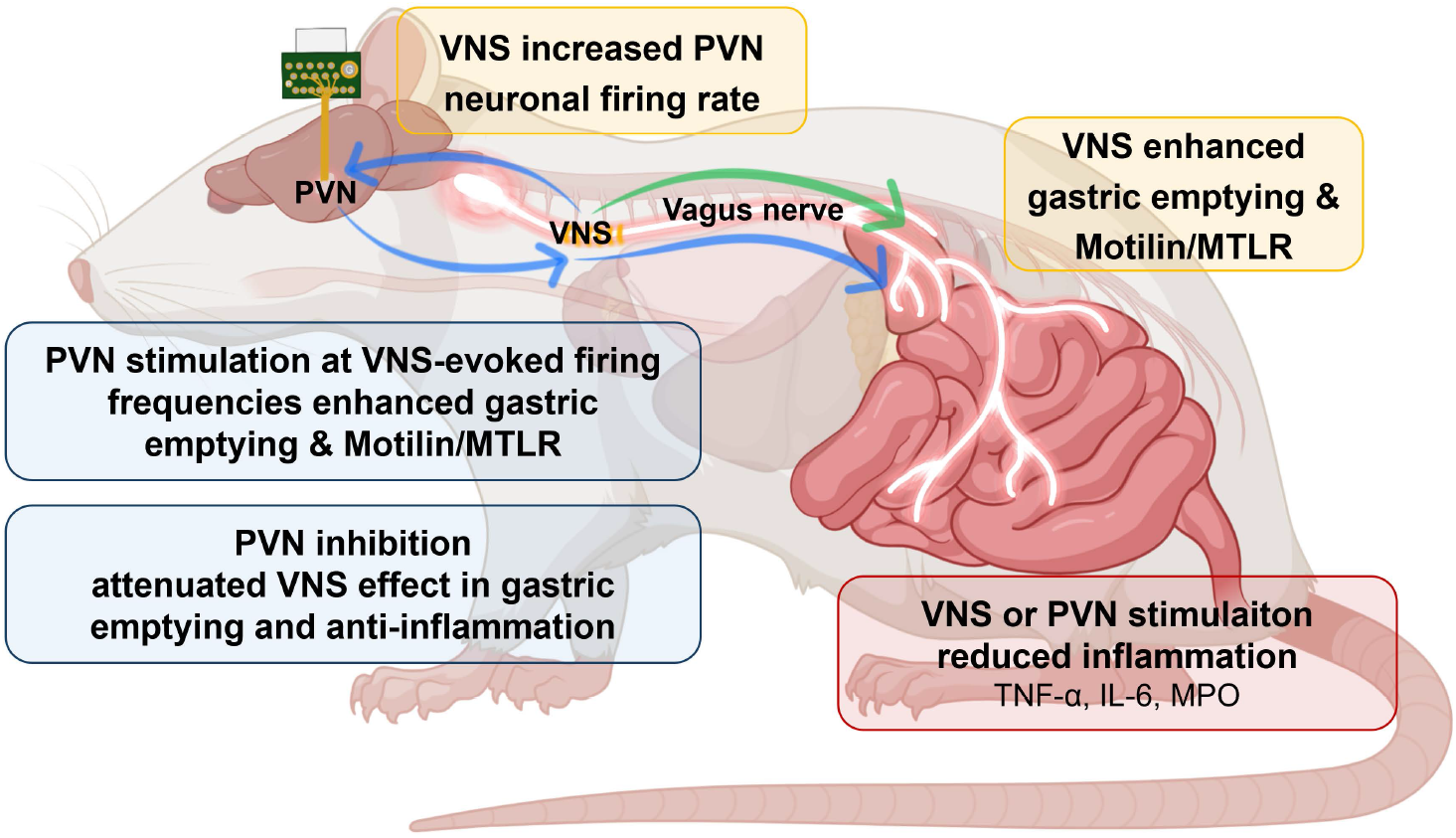

## 1. Introduction

Gastroparesis is a gastrointestinal (GI) motility disorder that is characterized by delayed gastric emptying (GE). This disorder may develop as a secondary complication of diabetes mellitus, abdominal surgery, medication use, autoimmune disorders, or neurological diseases and can impair nutritional status and reduce the quality of life [1,2]. Current treatments for gastroparesis rely primarily on prokinetic agents, which usually have limited efficacy or intolerable adverse effects [3]. Thus, there remains an unmet clinical need for mechanism-based neuromodulatory therapies targeting GI motility impairment. Within the GI system, vagal pathways regulate gastric accommodation, pyloric function, gastrointestinal transit, and GE through vagovagal reflexes [4,5]. Vagal afferents convey visceral information to the nucleus tractus solitarius (NTS), which then modulates neurons in the dorsal motor nucleus of the vagus (DMV) through inhibitory GABAergic and excitatory glutamatergic synaptic pathways. Preganglionic neurons of the DVM then provide vagal efferent output to enteric neurons that regulate GI motor function [6].

Vagus nerve stimulation (VNS) is a neuromodulation therapy approved by the Food and Drug Administration for drug-resistant epilepsy, treatment-resistant depression, and headache disorders [7]. Evidence suggests that VNS can modulate autonomic, gastrointestinal, and immune functions through bidirectional vagal signaling [8], and it facilitates digestion primarily by driving gastric and intestinal motility via vagal efferent synapses within the enteric nervous system [9]. This is achieved through the gastric excitatory vagal motor circuit that releases acetylcholine to trigger coordinated peristalsis and accelerate GE and intestinal transit [10] while the parallel gastric inhibitory circuits utilize nitric oxide and vasoactive intestinal peptide (VIP) to induce receptive relaxation of the stomach wall and sphincters for smooth food passage [11]. VNS also induces gastric acid and pancreatic enzyme secretions via muscarinic and VIP pathways for nutrient breakdown and local vasodilation to increase mesenteric blood flow for nutrient absorption [12].

In addition to autonomic regulation of gastric motility and digestion, VNS exerts potent neuro-immunomodulatory effects, suggesting therapeutic potential for inflammatory GI disorders [13]. One established mechanism is the vagal efferent cholinergic anti-inflammatory pathway (CAP) [6,14], through which VNS restores inflammatory related GI disorders [15]. Another proposed neuroimmune mechanism of VNS involves hypothalamic–pituitary–adrenal (HPA) axis activation, wherein vagal afferent signaling activates hypothalamic neurons, stimulating corticotropin-releasing hormone (CRH) and downstream glucocorticoid release [16]. Motilin is a key GI hormone that regulates gastric motility by sequentially activating motor complexes [17]. It is secreted primarily by enteroendocrine M cells in the proximal small intestine and exerts prokinetic effects through motilin receptors (MTLR) in GI smooth muscle and enteric neurons [18]. Clinically, erythromycin-mediated motilin receptor activation has demonstrated therapeutic benefit for gastroparesis, supporting a causal role of motilin signaling in GE [19].

The paraventricular nucleus (PVN) of hypothalamus is a major integrative center for autonomic and neuroendocrine regulation and a plausible mediator linking vagal afferent input to GI motor function [20]. PVN receives projections from NTS and sends descending projections to autonomic brainstem nuclei, including DMV [21–23]. PVN contains heterogeneous neurons, including neuroendocrine and pre-autonomic neurons, enabling coordination of stress responses, inflammation, feeding behavior, and visceral homeostasis [24,25]. PVN activity has been implicated in the regulation of gastric accommodation and distension and feeding behavior [5,26]. However, the role of PVN in regulating gastric motility and the prokinetic therapeutic effects of VNS remain to be understood.

It is crucial but less known how brain regulates VNS effects. The brain processes VNS afferents through a bottom-up polysynaptic pathway [27]. VNS signals propagated via vagal afferents directly to the NTS and then to the locus coeruleus and dorsal raphe nucleus, leading to norepinephrine (NE), serotonin (5-HT), and acetylcholine (ACh) release across the cortex, hippocampus, and amygdala. VNS desynchronizes aberrant cortical activity to suppress seizures, enhance synaptic plasticity for stroke rehabilitation, and downregulate neuroinflammation [13]. However, the top-down brain modulation of VNS peripheral effects remains largely unexplored. The PVN receives ascending projections from NTS, a major central relay for VNS-driven vagal afferent input. We therefore hypothesized that cervical VNS activates the PVN via this afferent route, where the PVN acts as a central regulator gating the VNS-enhanced descending vagal efferent output to the upper GI tract. Concurrently, this pathway is mediated through parallel vagovagal neural control and vagal neuroimmunomodulation. To investigate whether VNS improves gastric motility through a PVN-dependent mechanism, we examined PVN neuronal activity during VNS; the effects of PVN stimulation and PVN inhibition during VNS on GE in a rat model of gastroparesis; and motilin, corticosterone, and inflammatory cytokines as candidate downstream mediators.

## 2. Methods

### 2.1. Animal and intestinal manipulation model

Male Sprague–Dawley rats (8–10 weeks old, 250–350 g, BioLASCO, Taiwan) were housed under standard laboratory conditions with ad libitum access to food and water. All animal procedures were approved by the Institutional Animal Care and Use Committee of National Cheng Kung University (IACUC No. 114044). The animals were randomly divided into an intestinal manipulation (IM)-induced gastroparesis group and a control laparotomy (LP) group. Following a 20-h fast, rats were anesthetized with isoflurane (4%–5% for induction; 1%–3% for maintenance via nose cone) and subjected to abdominal midline incision in a supine position. Gastroparesis was induced using a modified IM model [28], wherein the GI tract was exteriorized and mechanically manipulated with sterile saline-moistened cotton swabs from the pyloric antrum to cecum for 5 min using a standardized force of 130–150 g (Supplementary Fig. 1). In the control LP group, the GI tract was gently exteriorized but then returned to the abdominal cavity without manipulation. The abdominal wall was subsequently closed.

### 2.2. GE and GI transit

GE and GI transit were quantified using fluorescein isothiocyanate (FITC)–dextran distribution analysis. Immediately after undergoing LP or IM, rats underwent oral gavage with 1.5 mL of FITC–dextran (Sigma-Aldrich, USA). After 3 hours, 6 mL of blood was collected via cardiac puncture under anesthesia. Thereafter, the GI tract or brain was harvested after transcardial perfusion with 4% paraformaldehyde (PFA). The excised whole GI tract was divided into 15 sequential segments comprising the stomach, 10 equal-length segments of the small intestine (sb1–sb10), the whole cecum, and three colonic segments (col1–col3; Fig. 1A). Luminal FITC–dextran was recovered and fluorescence at excitation and emission wavelengths of 485 and 528 nm, respectively, were measured (Supplementary Methods 1). GI transit was expressed as the FITC–dextran fluorescence intensity distributed over the GI tract. GE was calculated as GE (%) = [1 − (gastric fluorescence/total GI tract fluorescence)] × 100. Geometric center (GC) was calculated as the weighted average of the FITC–dextran position within the GI tract as follows: GC = Σ (% fluorescence per segment × segment number)/100. A lower GC, corresponding to a more proximal distribution of FITC–dextran, indicated delayed GI transit [29].

**Fig. 1.**
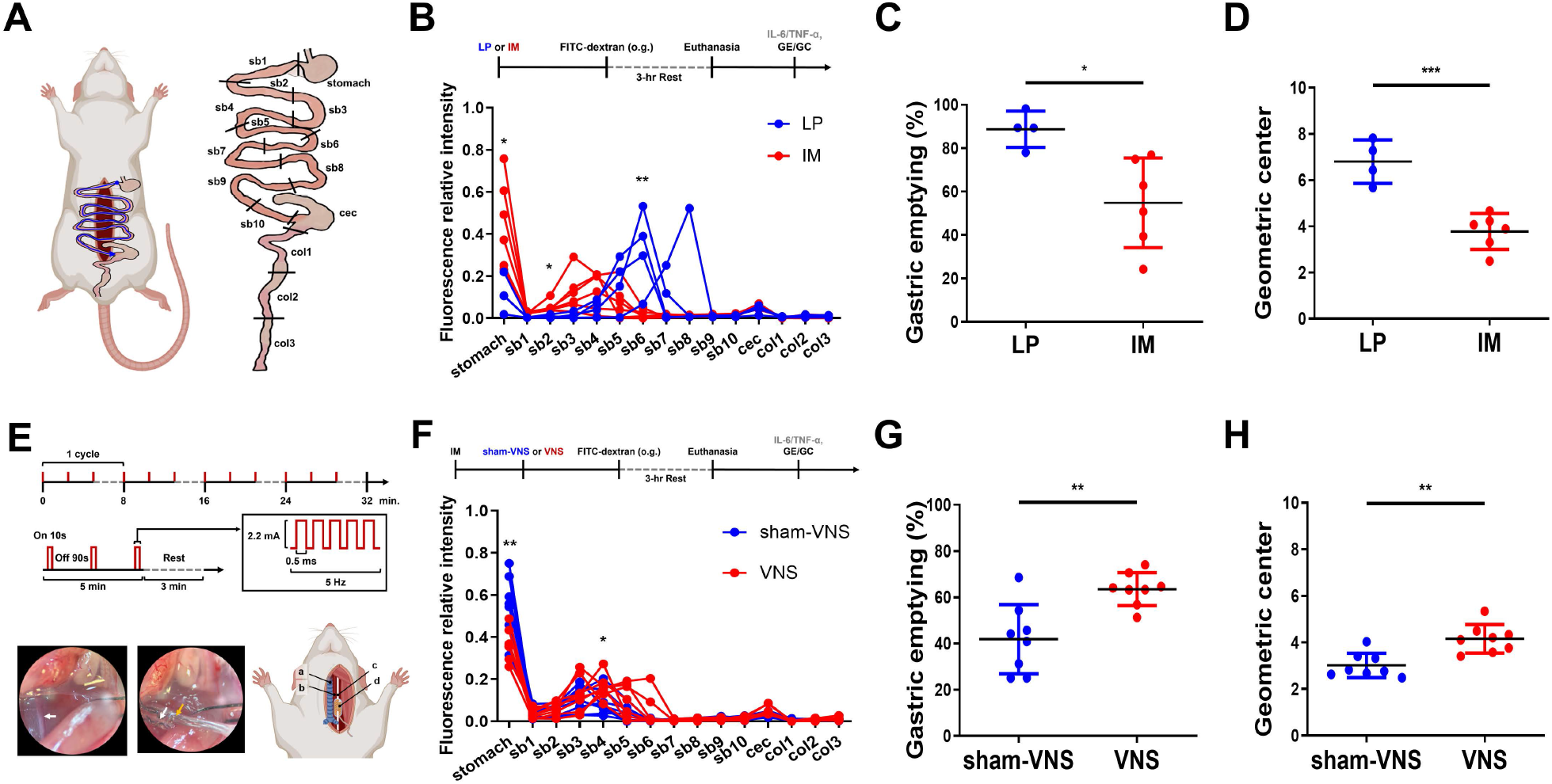
Vagus nerve stimulation (VNS) enhances gastrointestinal transit and gastric emptying in rats with intestinal manipulation (IM)-induced gastroparesis. **A,** Establishment of the IM-induced gastroparesis rat model and segmentation of the GI tract for FITC–dextran distribution measurement. **B,** Experimental protocol and distribution of FITC–dextran along the whole GI tract of gastroparesis rats (n = 6) compared with laparotomy (LP) controls (n = 4). FITC–dextran intensity, IM vs. LP: 0.45 ± 0.20 vs. 0.11± 0.08 at stomach; 0.05 ± 0.02 vs. 0.008 ± 0.007 at sb2; 0.016 ± 0.01 vs. 0.32 ± 0.195 at sb6. Each line represents an individual rat. **C,** Gastric emptying in gastroparesis rats (54.8 ± 20.7%) compared with LP controls (88.7 ± 8.3%). **D,** Geometric center of FITC–dextran distribution in gastroparesis rats (3.8 ± 0.8) compared with LP controls (6.8 ± 0.9). **E,** VNS protocol (upper panel) and electrode implantation (lower panels). Surgical microscope images show the exposed left cervical vagus nerve (white arrow, lower left) and the cuff electrode secured on the cervical vagus nerve (yellow and white arrows, respectively; lower middle). The schematic of the surgical field identifies the trachea (a), carotid artery (b), vagus nerve (c), and VNS cuff electrode (d). **F,** Experimental protocol and distribution of FITC–dextran along the whole GI tract from VNS-treated gastroparesis rats (n = 8) compared with the sham-stimulation rats (n = 8). FITC–dextran intensity, VNS vs. sham-stimulation: 0.36 ± 0.07 vs. 0.58 ± 0.14 at stomach; 0.16 ± 0.05 vs. 0.08 ± 0.05 at sb4. **G,** Comparison of gastric emptying between VNS (63.5 ± 7.1%) and sham-stimulation in IM-induced gastroparesis rats (41.9 ± 15.0%). **H,** Comparison of geometric center between VNS (4.2 ± 0.6) and sham-stimulation in IM-induced gastroparesis rats (3.0 ± 0.5, n = 8 per group). Data are presented as mean ± SD; n indicates the number of rats. *p < 0.05, **p < 0.01, ***p < 0.001, ****p < 0.0001. cec, cecum; col, colon; FITC, fluorescein isothiocyanate; GC, geometric center; GE, gastric emptying; IL-6, interleukin-6; o.g., oral gavage; sb, small bowel; TNF-α, tumor necrosis factor-α.

### 2.3. VNS

Under isoflurane anesthesia, the left cervical vagus nerve was gently isolated from the carotid sheath, and a nerve-cuff electrode (MicroProbes, USA) was positioned on it. Electrical contact confirmed by an acute decrease in heart rate during a brief test stimulation (Supplementary Fig. 2). To test GI motility enhancement, VNS was delivered at 2.2 mA and 5 Hz as 0.5-ms width square waves in 10-s stimulation/90-s rest cycles for 30 min (Fig. 1E). The sham-VNS group underwent identical procedures but without electrical stimulation. For measuring PVN response during VNS, VNS was delivered at 5 Hz as 0.5-ms width rectangular pulses with stepwise increases in current amplitude of 0.5, 1.0, 1.5, and 2.2 mA. Each stimulus was applied as 10-s stimulation/90-s rest cycles for 5 minutes, with 3-min resting intervals between different intensities (Fig. 2G).

**Fig. 2.**
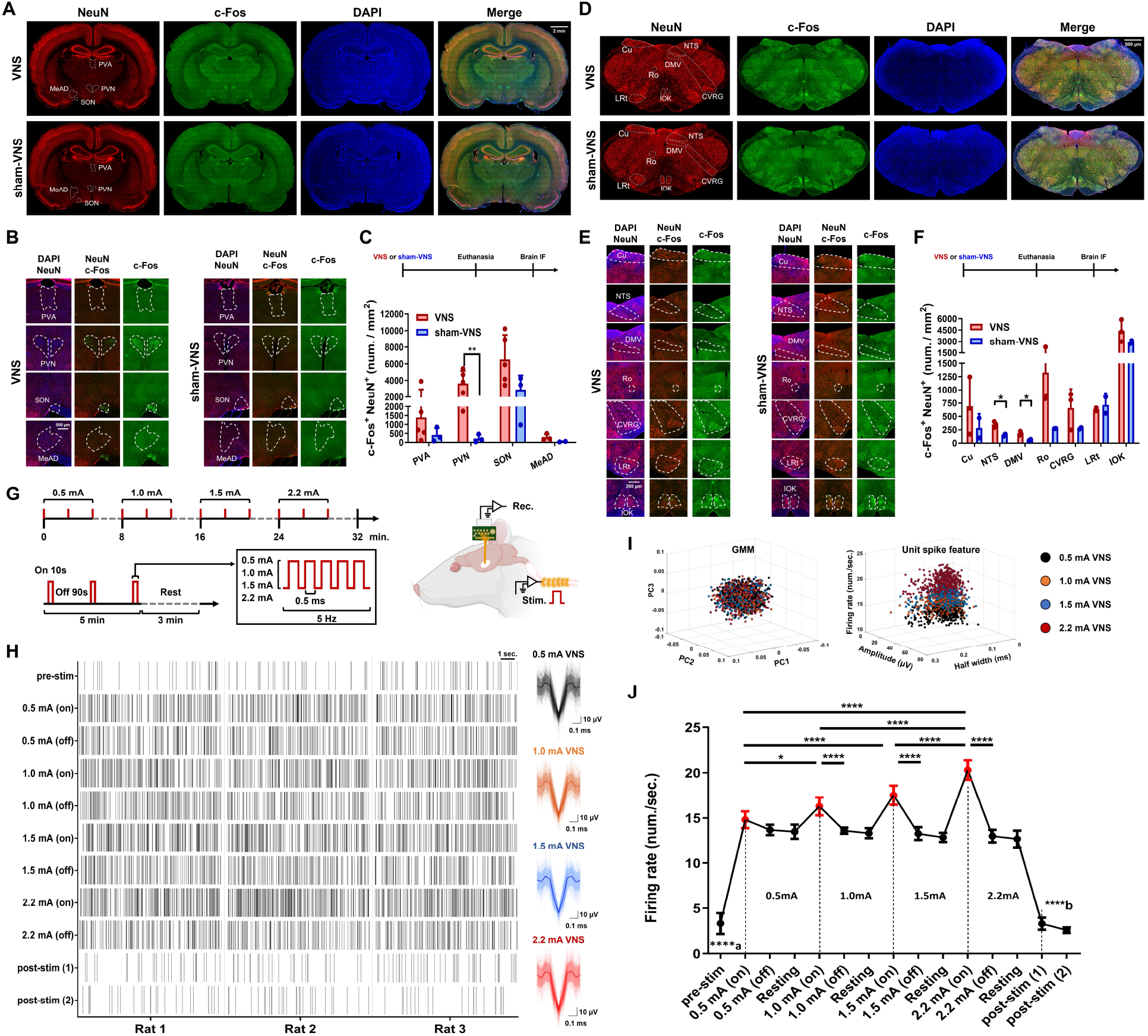
VNS activates the paraventricular nucleus (PVN), nucleus of the solitary tract (NTS), and dorsal motor nucleus of the vagus (DMV), and increases PVN neuronal firing in proportion to VNS intensity. A, Immunofluorescent images of the neuronal marker NeuN, neuronal activation marker c-Fos, nuclear marker DAPI, and merged images showing coexpression in whole-brain coronal sections containing PVA (paraventricular thalamic nucleus, anterior part), PVN, supraoptic nucleus (SON), and medial amygdaloid nucleus, anterodorsal part (MeAD) (dashed lines) from VNS-treated and sham-stimulated groups. **B,** Enlarged microscopic images with colocalized NeuN and DAPI (left), colocalized NeuN and c-Fos (middle), and c-Fos (right) within the PVA, PVN, SON, and MeAD of VNS-treated and sham-stimulated animals. **C,** Experimental protocol and comparison of neuronal activation indicated by c-Fos^+^ NeuN^+^ colocalized number between the VNS-treated group (n = 5, 12 slices) and sham-stimulated group (n = 3, 9 slices). c-Fos^+^ NeuN^+^ colocalized number at PVN, VNS: 3631.8 ± 1515.2 /mm² vs. sham VNS: 214.2 ± 220.6 /mm². **D,** Immunofluorescent images of NeuN, c-Fos, DAPI, and merged images of coexpression within whole-brain coronal sections containing the cuneate fasciculus (Cu), NTS, DMV, nucleus of Roller (Ro), caudoventral respiratory group (CVRG), lateral reticular nucleus (LRt), and inferior olive, cap of Kooy of the medial nucleus (IOK) (dashed lines) from VNS-treated and sham-stimulated rats. **E**, Enlarged microscopic images with colocalized NeuN and DAPI (left), colocalized NeuN and c-Fos (middle), and c-Fos (right) within the Cu, NTS, DMV, Ro, CVRG, LRt, and IOK from VNS-treated and sham-stimulated groups. **F,** Experimental protocol and comparison of the c-Fos^+^ NeuN^+^ neuron number between the VNS-treated group (n = 3, 9 slices) and sham-stimulated group (n = 2, 6 slices). c-Fos^+^ NeuN^+^ colocalized number at NTS, VNS: 349.6 ± 55 /mm² vs. sham VNS: 149.1 ± 28.7 /mm²; DMV, VNS: 181.6 ± 38.5 /mm² vs. sham VNS: 59.5 ± 15.9 /mm². **G,** VNS stimulation protocol (left) and simultaneous extracellular unit spike recording from PVN during VNS (right). **H,** Example raster plots of PVN unit spikes from a 10-s segment of each period (pre-, during, and post-VNS). Spike waveform recorded under each stimulation intensity presented with mean and SD indicated by line with shadow (n = 396, 444, 446, and 448 unit spikes, respectively,). **I,** Gaussian mixture model (GMM) clustering of PVN unit spikes during VNS (left) and waveform features of firing rate, half width, and amplitude (right). **J,** Comparisons of PVN neuronal firing rates during pre-stim, post-stim, VNS on, VNS off, and resting periods with varying current strength (n = 3). **a**, pre-stim compared with each VNS-on condition (0.5, 1.0, 1.5, 2.2 mA), all p < 0.0001. **b**, post-stim (1), the first hour after stimulation and post-stim (2), the second hour after stimulation compared with each VNS-on condition (0.5, 1.0, 1.5, 2.2 mA), all p < 0.0001. Data are presented as mean ± SD; n indicates the number of rats. *p < 0.05, **p < 0.01, ***p < 0.001, ****p < 0.0001. IF, immunofluorescence staining; rec., recording; stim., stimulation.

### 2.4. PVN recording, electrical stimulation, and pharmacological inhibition

Rats were positioned in a stereotaxic apparatus under isoflurane anesthesia and custom-fabricated tungsten-wire electrodes (50.8-μm diameter, HML-insulated; California Fine Wire Company, Grover Beach, CA, USA) were implanted into bilateral PVN (AP −1.8 mm, ML ±0.5 mm, DV −7.8 mm) with channels 1–4 and 5–8 at the left and right sides, respectively, linked to a connector (SenonLink Technology Co., Taiwan). Extracellular PVN neuronal activity was recorded bilaterally using a multichannel acquisition system (Plexon, USA) at a 20-kHz sampling rate with band-pass filtering at 300–6000 Hz and analyzed offline using UltraMegaSort2000 [30] and MATLAB for unit spike sorting and firing frequency [31] (Supplementary Methods 2). For direct PVN stimulation, channels 1–2 at left PVN delivered electrical current with 20 μA as 20-Hz sinusoidal waves in 10-s stimulation/90-s rest cycles for 30 min (Fig. 3A). For pharmacological inhibition, 70 nL of normal saline containing GABA_A_ agonist muscimol (0.5 nmol, Sigma-Aldrich, USA) was injected into bilateral PVN at AP –1.8 mm, ML ±0.5 mm, DV –7.8 mm using a calibrated glass micropipette through a pressure-driven microinjection system (Longer Pump, China) at a controlled infusion rate of 50 nL/2 min. The control group was injected with vehicle saline using the same procedures.

**Fig. 3.**
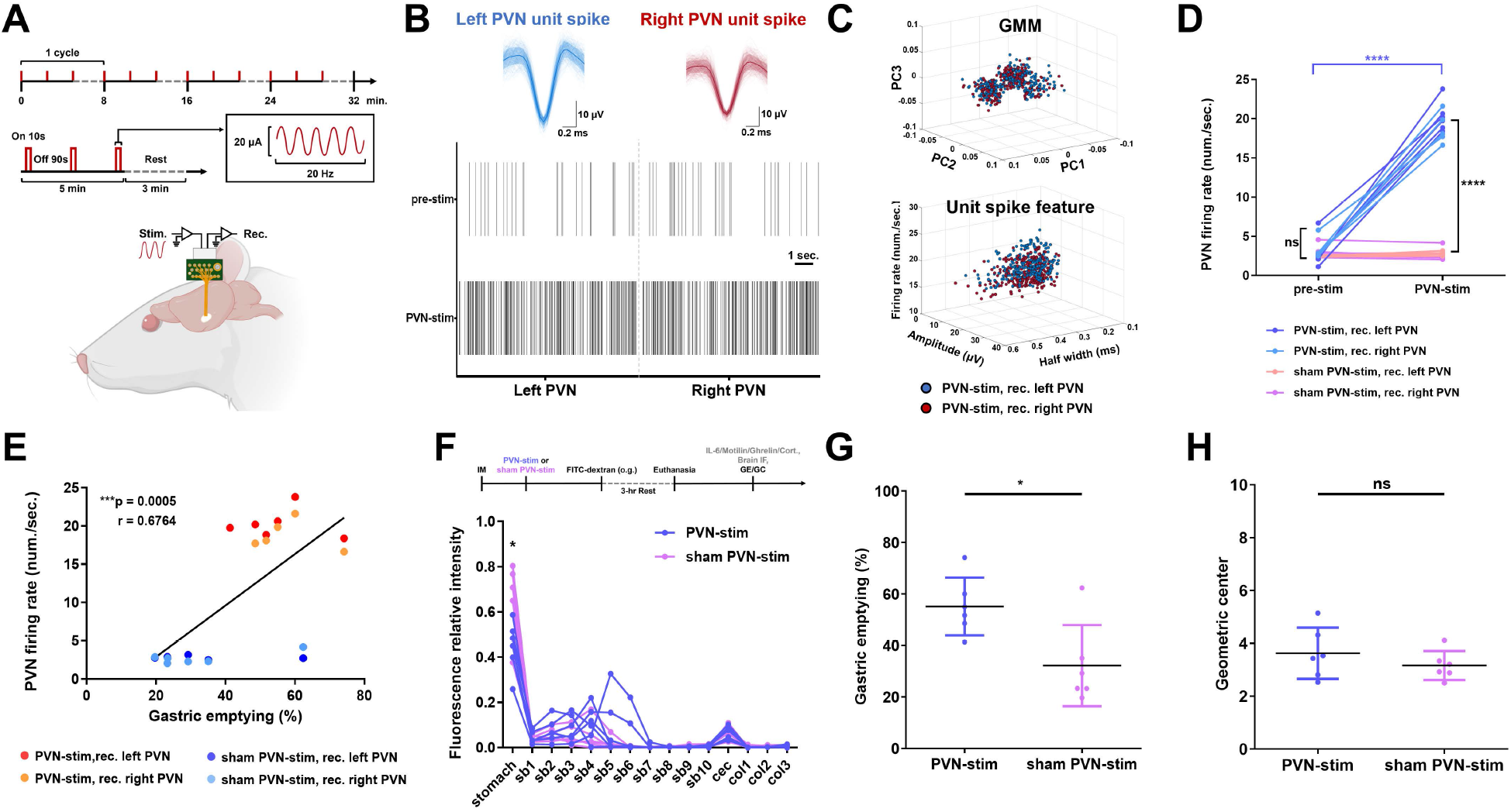
Direct electrical stimulation of PVN at VNS-evoked firing frequency improves gastric emptying in IM-induced gastroparesis rats. **A,** PVN stimulation protocol (upper) and simultaneous extracellular unit spike recording at bilateral PVNs during left PVN stimulation (lower). **B,** Representative raster plots in 10-s segment of PVN unit spikes before, and during left PVN stimulation. Waveform of unit spikes presented with mean and SD indicated by line with shadow (n = 262 and 305 unit spikes from left and right PVN, respectively). **C,** Gaussian mixture model (GMM) unit spike clustering and waveform features of PVN neurons during left PNV stimulation. Blue dots indicate unit spikes recorded from left PVN, red dots from right PVN. **D,** Comparison of PVN unit spike firing frequency between pre-stimulus and during left PVN stimulation (n = 6) or sham stimulation (n = 6). **E,** Correlation between PVN neuronal firing rate and gastric emptying among IM-induced gastroparesis rats receiving PVN stimulation or sham stimulation. **F,** Experimental protocol and comparison of FITC–dextran distribution along the whole GI tract of gastroparesis rats receiving PVN stimulation or sham stimulation. PVN stimulation: 0.44 ± 0.11 vs. sham stimulation: 0.67 ± 0.15 at stomach. Each line represents an individual rat. **G**, Comparison of gastric emptying between gastroparesis rats receiving PVN stimulation (55.1 ± 4.6%) and sham stimulation (32.2 ± 6.4%, n = 6 per group). **H**, Comparisons of FITC–dextran geometric center between gastroparesis rats receiving PVN stimulation (3.6 ± 0.9) and sham stimulation (3.6 ± 1, n = 6 per group). Data are presented as mean ± SD; n indicates the number of rats. *p < 0.05, **p < 0.01, ***p < 0.001, ****p < 0.000, ns, non-significant. Cort., corticosterone; GC, geometric center; GE, gastric emptying; IF, immunofluorescence staining; IL-6, interleukin-6; o.g., oral gavage; rec., recording; stim., stimulation.

### 2.5. Biochemical and histological analyses

Serum interleukin-6 (IL-6), tumor necrosis factor-α (TNF-α), motilin, ghrelin, and corticosterone were measured using commercial ELISA kits. Following transcardial perfusion with saline and 4% PFA, brains were post-fixed overnight at 4°C, cryoprotected in 30% sucrose, and coronally sectioned at 40 μm. Whole-brain sections were immunostained for NeuN, DAPI, and c-Fos, and PVN-containing sections were immunolabeled for CRH and c-Fos, with colocalization assessed in the respective experiments. GI tissues were paraffin-embedded and immunostained for MTLR or myeloperoxidase (MPO), a leukocyte marker, in respective experiments. Images were acquired using a confocal laser scanning microscope (FV3000, Olympus) and quantified using a panoramic tissue cytometry system (TissueFAXS, TissueGnostics) with TissueQuest software. Antibody information, staining and quantification procedures, and material details are provided in Supplementary Methods 3 and Supplementary Table 1.

### 2.6. Statistical analysis

Data are presented as mean ± SD from n rats. Two-group comparisons used unpaired t-tests for normally distributed data and Welch’s t-test for unequal variances. Repeated measurements used one-way ANOVA with post-hoc Bonferroni’s tests. Association were assessed using Pearson’s correlation coefficient. GraphPad Prism 11 were used; P <0.05 indicated statistically significant.

## 3. Results

### 3.1. VNS enhances GI transit and GE in IM-induced gastroparesis rats

Gastroparesis was successfully induced in rats subjected to IM, as evidenced by significantly reduced GE and shorter the FITC–dextran transit distance compared with control LP rats, with greater FITC–dextran retention in the stomach and proximal GI tract (p = 0.015 for stomach; p = 0.012 for sb2; p = 0.004 for sb6; unpaired t-test; Fig. 1A and 1B). Compared with LP controls, IM group exhibited significantly reduced GE (p = 0.015; Fig 1C) as well as GC (p = 0.0005; Fig. 1D; unpaired t-test). Application of a 2.2-mA VNS cycling protocol for 30 min ameliorated gastroparesis in IM-treated rats (Fig. 1E), as evidenced by increased GE and greater FITC– dextran transit distance compared with sham-stimulation, with reduced FITC–dextran retention in the stomach (p = 0.002) and increased in sb4 (p = 0.0228, unpaired t-test; Fig. 1F). Compared with sham-IM group, GE and GC improved in the VNS-IM group (p = 0.002 at GE, Fig 1G; p = 0.001 at GC, Fig 1H; unpaired t-test).

### 3.2. VNS activates PVN, NTS, and DMV and increases PVN neuronal firing in a stimulation-strength dependent response

To investigate brain activation by VNS, whole-brain sections, including brainstem nuclei were examined by c-Fos immunostaining following VNS (Fig. 2A–2F). Compared with sham-stimulation group, the VNS group showed significantly enhanced c-Fos neuronal activation at PVN (p = 0.007; Fig. 2C), NTS (p = 0.013), and DMV (p = 0.019; Fig. 2F, by Welch’s t-test,). The unit spikes measured from PVN were sorted into one cluster with consistent waveforms across varied stimulation intensity with progressively greater frequencies at higher VNS intensities (Fig. 2G–2I; Supplementary Table 2). Specifically, the mean firing frequency of PVN neurons significantly increased as VNS intensity increased in a stepwise manner compared with the pre- and off-stimulation and resting state of each session (stimulation condition, F_(14, 120)_ = 385.2, p < 0.0001, one-way ANOVA; Fig. 2J; post-hoc test, Supplementary Table 3). The PVN firing rate increased from a basal rate of 3.3/s to 14.8, 16.3, 17.5, 20.3/sec at 0.5, 1.0, 1.5, and 2.2 mA VNS, respectively (p < 0.0001 at each stimulation strength compared with pre-stimulation; Fig 2J); it returned to 3.3/s after stimulation. VNS at 2.2 mA significantly improved GI motility (Figure 1). These findings support the presence of a functional vagal-to-PVN pathway where PVN neurons were activated by VNS.

### 3.3. Direct electrical stimulation of PVN at the firing frequency analogous to VNS-induced responses improves GE in MI-induced gastroparesis rats

We further test the causality of PVN in GI motility by applying 20 µA sinusoid electrical stimulation at 20 Hz to the left PVN in analogue to its firing frequency during 2.2 mA VNS (Fig. 3A). Unit spikes exhibited a consistent waveform and features in bilateral PVN (Fig. 3B and 3C, Supplementary Table 2) and a significant increase in frequency from 3.1 ± 1.6/s at baseline to 19.6 ± 2/s during stimulation (p < 0.0001, unpaired t-test; Fig. 3D). Importantly, the PVN firing frequency was positively correlated with GE (r = 0.676, p = 0.0005, Pearson’s correlation test; Fig. 3E). The distribution of FITC–dextran in the stomach reduced in PVN-stimulation group compared with sham-stimulation group (p = 0.016 at stomach, unpaired t-test; Fig. 3F). In IM-induced gastroparesis rats, GE significantly increased in PVN-stimulation group compared with sham-stimulation group (p = 0.016, unpaired t-test; Fig. 3G). However, unlike VNS, GC did not improve in the PVN-stimulation group (p = 0.325, unpaired t-test; Fig. 3H) showing a discrepancy of PVN regulation in gastric and intestinal motility.

### 3.4. Muscimol-induced PVN inhibition abolishes the prokinetic effects of VNS in IM-induced gastroparesis rats

To identify the role of PVN in the prokinetic effects of VNS, GE and GC were measured following pharmacological PVN blockade in VNS-treated IM-induced gastroparesis rats. c-Fos immunostaining (Fig. 4A) showed that direct PVN stimulation increased PVN neuronal activation versus sham-stimulation (p = 0.04), whereas PVN microinfusion of the GABA_A_ agonist muscimol markedly reduced VNS-induced activation versus saline-injection control (p = 0.002, Fig. 4B). Muscimol also significantly reduced GI transit with more FITC–dextran retention in stomach (p = 0.009) and proximal GI tract (p = 0.044 at sb3; p = 0.022 at sb5, unpaired t-test; Fig. 4C). In VNS-treated IM rats, PVN muscimol infusion reduced GE compared with saline-treated PVN controls (p = 0.009) and the VNS-only group (p = 0.009), to a level comparable to sham-VNS group (p = 0.497, unpaired t-test), indicating that VNS-induced GE enhancement was almost abolished by PVN blockade. In gastroparesis animals, GE did not differ significantly between those receiving PVN saline injection combined with VNS and those receiving VNS alone (p = 0.329; Fig. 4D). Similarly, GC reduced by muscimol compared with saline controls (p = 0.012, unpaired t-test; Fig. 4E), to a comparable level of sham-VNS group (p = 0.078; Fig. 4E). Results of the VNS- and sham-stimulation groups shown in Figures 4D and 4E are reproduced from Figures 1G and 1H for comparison. These findings support a critical role for PVN in VNS-mediated prokinetic effects and PVN-to-vagal regulation of upper GI motility. Post-procedure general activity did not differ among groups, excluding motor activity as a potential confounder (Supplementary Fig. 3).

**Fig. 4.**
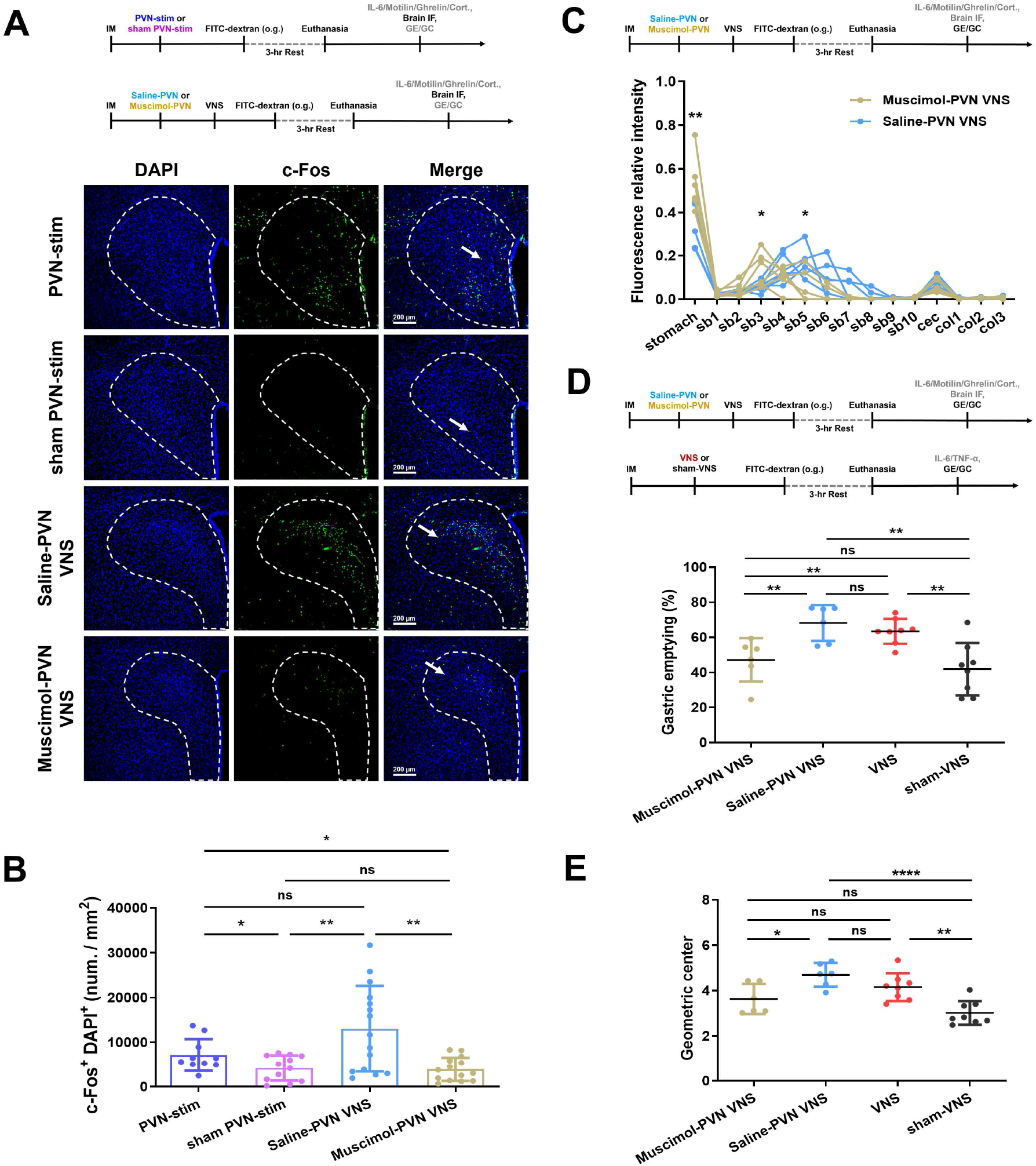
Muscimol inhibition of PVN abolishes the prokinetic effects of VNS in IM-induced gastroparesis rats. **A,** Experimental protocols and immunofluorescent images of DAPI (left), c-Fos (middle), and merged images (right) from PVN. Arrow indicates c-Fos^+^ cell. **B,** Comparisons of the neuronal activation labeled with c-Fos^+^DAPI^+^ within PVN in gastroparesis rats receiving PVN stimulation (7138 ± 3588/mm^2^, n = 6, 10 slices), PVN sham stimulation (4190 ± 2832/mm^2^, n = 6, 12 slices), VNS with saline injection into PVN (13005 ± 9605/mm^2^, n = 6, 15 slices) or VNS with muscimol injection into PVN (3895 ± 2634/mm^2^, n = 6, 14 slices). **C,** Comparison of whole GI FITC– dextran distribution between PVN muscimol-injected and PVN saline-injected gastroparesis rats receiving VNS (n = 6 rats per group). FITC–dextran intensity, muscimol vs. saline: 0.52 ± 0.12 vs. 0.31 ± 0.10 at stomach; 0.13 ± 0.07 vs. 0.05 ± 0.02 at sb3; 0.05 ± 0.07 vs. 0.16 ± 0.06 at sb5, Each line represents an individual rat. **D,** Experimental protocols and comparison of gastric emptying among gastroparesis rats receiving VNS with PVN saline injection (68.2 ± 10.3%, n = 6), VNS with PVN muscimol injection (47.1 ± 12.4, n = 6), VNS (63.5 ± 7.1%, n = 8), or sham-VNS stimulation (41.9 ± 15%, n = 8). **E,** Comparison of FITC–dextran geometric center among gastroparesis rats receiving VNS with PVN saline injection (4.7 ± 0.2, n = 6), VNS with PVN muscimol injection (3.6 ± 0.7, n = 6), VNS (4.2 ± 0.6, n = 8), or sham-VNS (3 ± 0.5, n = 8). Data are presented as mean ± SD; n indicates the number of rats. *p < 0.05, **p < 0.01, ***p < 0.001, ****p < 0.000, ns, non-significant. Cort., corticosterone; GC, geometric center; GE, gastric emptying; IF, immunofluorescence staining; IL-6, interleukin-6; o.g., oral gavage.

### 3.5. Enhanced GE in IM-induced gastroparesis rats after PVN stimulation and VNS associated with elevated serum motilin levels

We further investigated peripheral factors mediating the prokinetic effects of PVN stimulation and VNS on gastric motility. In gastroparesis rats, PVN stimulation increased serum motilin versus sham-stimulation (p < 0.0001), while VNS with saline injected PVN further increased motilin than PVN stimulation (p < 0.0001, unpaired t-test). However, PVN inhibition did not reduce motilin compared with saline injection following VNS (p = 0.707), potentially reflecting direct VNS efferent effects and a smaller contribution of PVN activation than VNS to motilin release. Notably, serum motilin levels were positively correlated with GE in rats receiving PVN stimulation or VNS (r = 0.437; p = 0.0124, Pearson’s correlation test; Fig. 5B). Gastric MTLR expression was increased after PVN stimulation versus sham-stimulation (p = 0.004), but not in the first small bowel segment (p = 0.623, unpaired t-test; Fig. 5C), consistent with the predominant effect of PVN stimulation on gastric rather than intestinal motility. In VNS-treated rats, PVN inhibition reduced MTLR expression compared with saline controls (stomach, p = 0.0003; sb3, p < 0.0001; sb5, p = 0.009; sb6, p = 0.0002, unpaired t-test; Fig. 5D). These findings suggest that enhanced motilin signaling may contribute to the prokinetic effects of PVN stimulation and VNS. Serum ghrelin was unchanged after PVN stimulation (p = 0.575) or PVN inhibition in VNS-treated rats (p = 0.274, unpaired t-test; Fig. 5E).

**Fig. 5.**
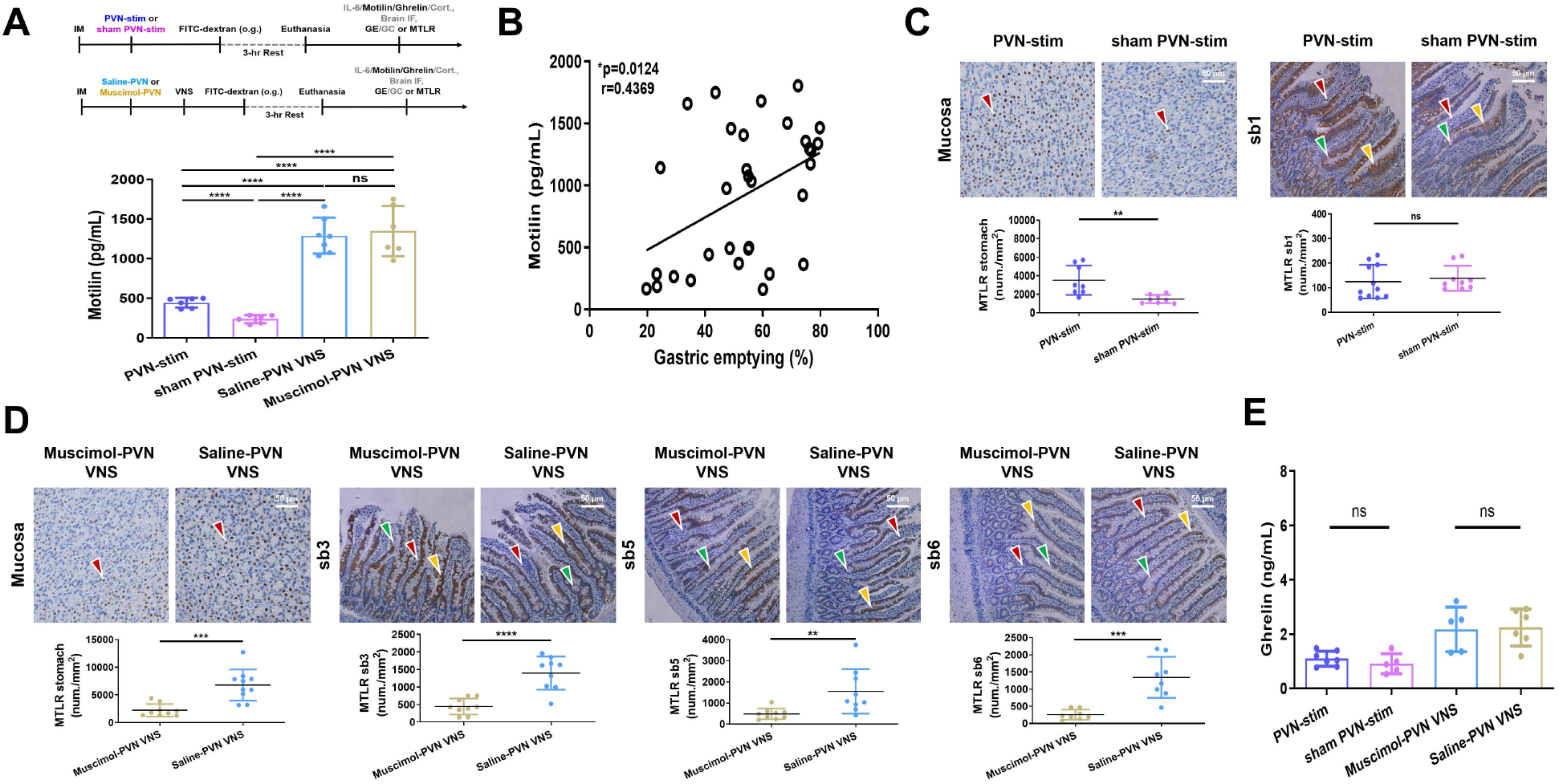
Enhanced gastric emptying in IM-induced gastroparesis rats after PVN stimulation and VNS associated with elevated serum motilin levels. **A,** Experimental protocols and comparison of serum motilin levels among gastroparesis rats receiving PVN stimulation (402.6 ± 62.3 pg/mL, n = 6), PVN sham stimulation (237.0 ± 51.4 pg/mL, n = 6), VNS with PVN muscimol injection (1347 ± 317.7 pg/mL, n = 6), or VNS with PVN saline injection (1289 ± 226.1 pg/mL, n = 7). **B,** Correlation between serum motilin concentration and gastric emptying among gastroparesis rats receiving PVN stimulation, PVN sham stimulation, VNS with PVN muscimol injection, or VNS with PVN saline injection. **C,** Comparison of motilin receptor (MTLR) postive cells in stomach (left, 3515 vs. 1479 /mm^2^, n = 8 slices per group) and sb1 (right, 124.9 vs. 138.7 /mm^2^, n = 11, 9 slices) between gastroparesis rats receiving PVN stimulation or PVN sham stimulation. MTLR indicated by red arrows, goblet cells by green arrows, and epithelial cells by yellow arrows in the upper panel. Statistical analyses shown in the lower panel. **D,** Comparison of MTLR-positive cell in stomach (2227 vs. 6785 /mm^2^, n = 9, 10 slices), sb3 (443.9 vs. 1397 /mm^2^, n = 9 slices per group), sb5 (484.5 vs. 1551 /mm^2^, n = 9 slices per group), and sb6 (257.1 vs. 1342, n = 8 slices per group) of gastroparesis rats receiving VNS with PVN muscimol injection or VNS with PVN saline injection. MTLR indicated by red arrows, goblet cells by green arrows, and epithelial cells by yellow arrows in upper panel. Statistical analysis shown in the lower panel. **E,** Comparison of serum ghrelin concentrations among gastroparesis rats receiving PVN stimulation, PVN sham stimulation, VNS with PVN muscimol injection, or VNS with PVN saline injection. sb1, sb3, sb5 and sb6 indicating small bowel segments 1, 3, 5, and 6. Data are presented as mean ± SD; n indicates the number of rats. *p < 0.05, **p < 0.01, ***p < 0.001, ****p < 0.000, ns, non-significant. Cort., corticosterone; GC, geometric center; GE, gastric emptying; IF, immunofluorescence staining; o.g., oral gavage.

### 3.6. VNS alleviates systemic and local inflammation in IM-induced gastroparesis rats

To investigate if neuro-immunomodulation contributes to the prokinetic effects of VNS, inflammatory profiles were examined. Serum levels of IL-6 and TNF-α were significantly elevated after IM compared with LP control (IL-6, p = 0.045; Fig 6A; TNF-α, p = 0.032; Fig 6B, unpaired t-test), indicating a systemic inflammation in IM-induced gastroparesis. Notably, compared with gastroparesis rats receiving sham-VNS, VNS reduced serum levels of IL-6 (p = 0.011; Fig. 6C) and TNF-α (p = 0.035; Fig. 6D, unpaired t-test). MPO is an enzyme stored in the primary neutrophil granules that is activated when neutrophils migrate into inflamed intestinal tissue [32]. In animals with IM-induced gastroparesis, VNS decreased MPO expression in sb1 compared with sham (p = 0.015, unpaired t-test; Fig 6E), consistent with a local anti-inflammatory effect in the upper GI tract.

**Fig. 6.**
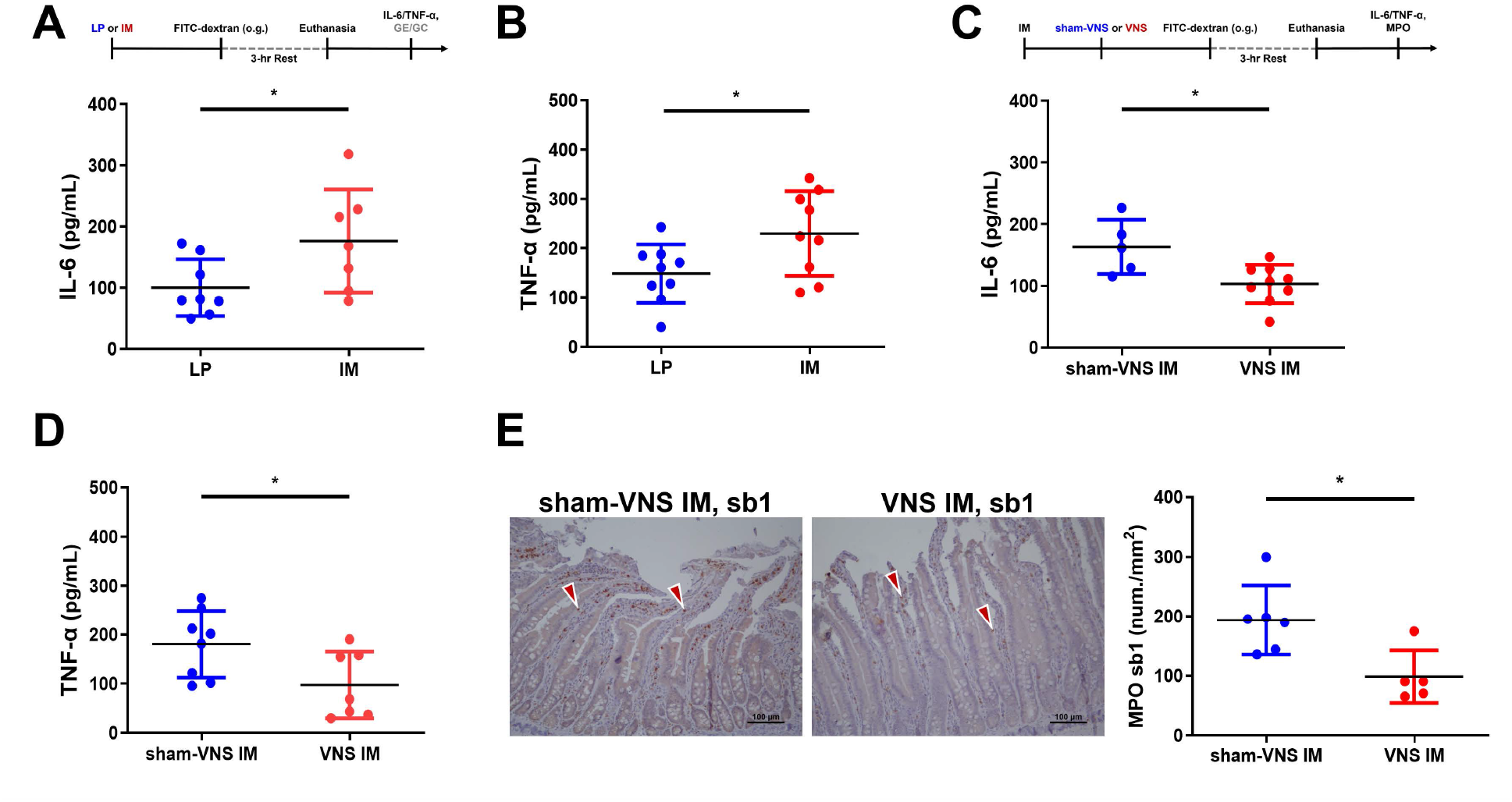
VNS alleviates systemic and local inflammation in IM-induced gastroparesis rats. **A**, Experimental protocol and comparison of serum IL-6 concentrations between IM-induced gastroparesis rats (176.4 pg/mL, n = 7) and LP controls (100.0 pg/mL, n = 8). **B**, Comparison of serum TNF-α concentrations between IM-induced gastroparesis rats (229.9 pg/mL, n = 9) and LP controls (148.3 pg/mL, n = 9). **C**, Experimental protocol and comparison of serum IL-6 concentrations between IM-induced gastroparesis rats receiving VNS (103.0 pg/mL, n = 9) and sham stimulation (163.2 pg/mL, n = 5). **D,** Comparison of serum TNF-α concentrations between IM-induced gastroparesis rats receiving VNS (97.21, n = 7) and sham stimulation (180.3 pg/mL, n = 8). **E**, Immunohistograph of MPO-positive cells (red arrows) in sb1 of gastroparesis rats receiving VNS or sham stimulation (left) and the comparison (right, 98.4, n = 5 vs. 193.7 /mm², n = 6). Data are presented as mean; n indicates the number of rats. *p < 0.05. GC, geometric center; GE, gastric emptying; IL-6, interleukin-6; MPO, myeloperoxidase; o.g., oral gavage; sb, small bowel; TNF-α, tumor necrosis factor-α.

### 3.7. PVN regulates the anti-inflammatory effects of VNS through reducing systemic IL-6 and local MPO expression in IM-induced gastroparesis rats

Finally, we examined whether PVN regulates VNS-enhanced immunomodulation. Serum IL-6 level was reduced after direct PVN stimulation versus PVN sham-stimulation (p = 0.045), whereas PVN inhibition with muscimol increased serum IL-6 versus saline control in VNS-treated rats (p = 0.041, unpaired t-test; Fig. 7A), indicating attenuation of the VNS-induced systemic anti-inflammatory effect. However, serum IL-6 level was not correlated with GE (r = 0.3096; p = 0.0846 by Pearson’s correlation; Fig. 7B). PVN stimulation also reduced MPO expression versus PVN sham-stimulation (p = 0.003 at sb1, p <0.0001 at sb4; Fig. 7C), whereas PVN inhibition increased MPO versus saline in VNS-treated gastroparetic model rats (p = 0.0002 at sb1, p = 0.0065 at sb4; Fig. 7D). CRH-positive neuronal activation increased after PVN stimulation versus sham-stimulation (p = 0.0012) and decreased following muscimol-induced PVN inhibition versus controls (p = 0.0117, unpaired t-test; Fig 7E). Serum corticosterone levels unchanged following PVN stimulation or PVN inhibition in VNS-treated gastroparesis rats compared with controls (Fig. 7F) and was not correlated with GE (r = -0.1987; p = 0.33; Pearson’s correlation; Fig. 7G).

**Fig. 7.**
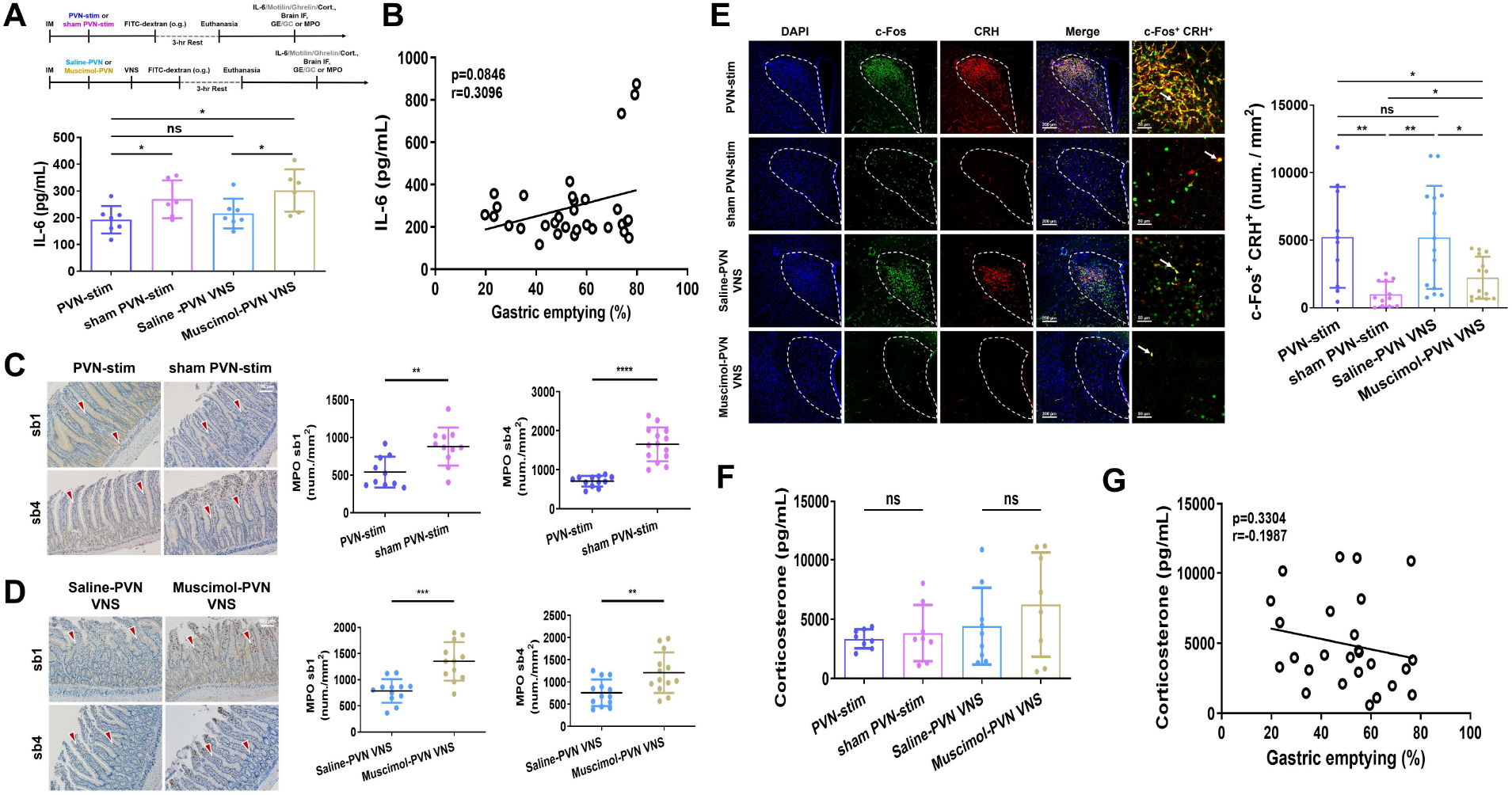
PVN regulates the anti-inflammatory effects of VNS through reducing systemic IL-6 and local MPO expression in IM-induced gastroparesis rats. **A,** Experimental protocols and comparison of serum IL-6 serum concentration among gastroparesis rats receiving PVN stimulation (192.5 ± 51.3 pg/mL, n = 7), PVN sham stimulation (269.0 ± 70.7 pg/mL, n = 6), VNS with PVN muscimol injection (301.8 ± 78.9 pg/mL, n = 6), or VNS with PVN saline injection (215.6 ± 55.3 pg/mL, n = 7). **B,** Correlation between serum IL-6 concentration and gastric emptying in gastroparesis rats receiving PVN stimulation, PVN sham stimulation, VNS with PVN muscimol injection, or VNS with PVN saline injection. **C,** Immunohistograph of MPO-positive cells (red arrows) in sb1 (540.7 ± 204 /mm², 10 slices vs. 880.8 ± 252.6 /mm², 11 slices) and sb4 (707.2 ± 136.2 /mm², 12 slices vs. 1650 ± 431.9 /mm², 14 slices) of gastroparesis rats receiving PVN stimulation or PVN sham stimulation (left) and comparison (right). **D,** Immunohistograph of MPO expression cells (red arrows) at sb1 (785.0 ± 227.3 /mm², 12 slices vs. 1353 ± 368.0 /mm², 12 slices) and sb4 (755.3 ± 298.9 /mm², 13 slices vs. 1208 ± 457.9 /mm², 13 slices) of VNS-treated gastroparesis rats receiving PVN muscimol injection or PVN saline injection (left) and comparison (right). **E,** Representative immunofluorescence images of PVN labeled with DAPI, c-Fos, CRH, and merged images. Enlarged images, experimental protocols and comparisons of c-Fos^+^CRH^+^ colocalization neurons at PVN, indicated by arrows, in gastroparesis rats receiving PVN stimulation (5210 ± 3735 /mm^2^, n = 10), PVN sham stimulation (1006 ± 924.6 /mm^2^, n = 12), VNS with PVN muscimol injection (2220 ± 1555 /mm^2^, n = 14), or VNS with PVN saline injection (5203 ± 3813 neurons/mm^2^, n = 14). **F,** Comparison of serum corticosterone among gastroparesis rats receiving PVN stimulation, PVN sham stimulation, VNS with PVN muscimol injection, or VNS with PVN saline injection. **G,** Correlation between serum corticosterone concentration and gastric emptying in IM-induced gastroparesis rats receiving PVN stimulation, PVN sham stimulation, VNS with PVN muscimol injection, or VNS with PVN saline injection. *p < 0.05, **p < 0.01, ***p < 0.001, ****p < 0.000, ns, non-significant. Cort., corticosterone; GC, geometric center; GE, gastric emptying; IF, immunofluorescence staining; IL-6, interleukin-6; o.g., oral gavage.

## 4. Discussion

The study demonstrates that cervical VNS improves GE in IM-induced gastroparesis rats through a vagal–PVN circuit, wherein PVN neurons activated by VNS afferents gate VNS efferent output to the GI tract and ensuing upregulation of motilin expressions. Both NTS and DMV can regulate upper GI motor functions, including gastric accommodation, antral-pyloric activity, and upper GI transit through vagovagal reflexes [23]. In addition to the vagovagal reflex involving vagus afferents, NTS, DMV, and vagal efferents, our findings reveal a crucial contribution of PVN to VNS-induced gastric motility improvement. First, NTS and DMV are well-established nuclei mediating vagal signals, and notably, our data revealed that PVN was activated by VNS to a greater extent than NTS and DMV as assessed by c-Fos expression. Besides, extracellular recordings demonstrated that PVN increased neuronal firing during VNS in a stimulation-strength dependent response, likely owing to the activation of a vagal afferent– NTS–PVN pathway. Second, direct PVN electrical stimulation significantly improved GE but not GC, with increased MTLR expression in the stomach but not intestine. These findings suggest that PVN stimulation preferentially enhances GE rather than intestinal transit, potentially supporting a more targeted approach for stomach motility disorders, whereas VNS improves both GE and intestinal motility. Serum motilin level was also lower under PVN stimulation than VNS, possibly due to differences in stimulation intensity and resultant activity of enteric motor neurons. Third, pharmacological PVN inhibition by local muscimol infusion attenuated the VNS-induced GE and GC enhancement, suggesting that intact PVN activity is necessary for gating the prokinetic effects of VNS. Collectively, these results support PVN as a critical regulator of GE and gating of upper GI kinetics enhanced by VNS. These findings also suggest that VNS enhances GE through peripheral and central pathways. Peripherally, VNS may directly activate vagal efferent pathways to the upper GI tract, thereby modulating gastric function. Centrally, we propose the involvement of a bidirectional vagal–PVN circuit that gates the VNS prokinetic effects. In this model, VNS afferents activate PVN, which in turn modulates descending vagal efferent output to the upper GI, potentially by engaging a vagal–NTS–PVN–DMV–vagal pathway [5].

This study also identified a positive correlation between motilin signaling and enhanced GE following PVN stimulation or VNS with PVN manipulation. Ghrelin was not significantly associated with GE. These findings support motilin as a potential peripheral mediator of the PVN-gated VNS prokinetic effect on GE. In contrast to the preferential gastric effect of PVN stimulation, VNS improved both GE and GC and induced higher motilin levels.

The IM-induced gastroparesis rat model is widely used to study GI dysmotility disorders, particularly postoperative delayed GE and inflammatory conditions, as well as the prokinetic effects of various potential treatments [33]. In our model, impaired GE and GC were accompanied by systemic and local GI inflammatory profiles characterized by high serum IL-6 and TNF-α levels and tissue MPO expression, a marker for neutrophil infiltration. Cervical VNS improved GE and GC while concomitantly reducing systemic and GI inflammation signs, consistent with previous reports [34,35]. This also supports VNS as a potential treatment for inflammatory GI motility disorders. Previous reports have suggested that neuroimmunomodulation by VNS can be mediated through peripheral CAP [13,36]. We further observed that PVN stimulation had similar anti-inflammatory effects, reducing both serum IL-6 and intestinal MPO levels, possibly by activating the same vagal neuroimmunomodulation pathway. Conversely, PVN inhibition abolished these VNS-induced anti-inflammatory effects, suggesting that the PVN may also gate the downstream anti-inflammatory actions of VNS. However, serum IL-6 level was not correlated with GE, suggesting that these anti-inflammatory effects do not directly or immediately mediate the gastric-specific prokinetic effects of VNS, at least in this acute model. Rather, inflammation suppression is likely to contribute to a broader recovery of gastrointestinal function in the subacute or chronic stage. Taken together, these findings suggest that VNS exerts its anti-inflammatory effects primarily through peripheral vagal neuroimmunomodulation, while the PVN may centrally regulate these effects via the vagal–PVN circuit.

PVN contains CRH-expressing neurons that regulate activation of the HPA axis and stress responses [5]. In the present study, although VNS and PVN stimulation activated CRH-positive PVN neurons, serum corticosterone remained unchanged and was not associated with GE. Stress-related CRH signaling is commonly associated with suppression of upper GI motility [37]. Our data do not support the role of the CRH–corticosterone signaling as a mediator of the prokinetic effects of VNS or PVN stimulation.

This study has several limitations. First, PVN contains heterogeneous neurons, including CRH and oxytocin neurons implicated in GE and GI inflammation, but the cell types activated by electrical stimulation were not identified [38,39]. Second, whether VNS–PVN activation regulates gastric motility independently of vagal efferents remains unclear. Third, given the lack of a functional motilin–receptor system in rats [17], the specificity of serum motilin and gastric MTLR measurements requires validation, and their causal involvement in VNS-enhanced gastric emptying remains uncertain. Finally, whether central PVN regulation of VNS effects is specific to upper GI motility or involves other autonomic functions remains unknown.

## 5. Conclusion

The present study suggests that a vagal–PVN–motilin pathway contributes to the prokinetic effects of VNS in IM-induced gastroparesis rats. Our findings further suggest a novel bidirectional vagal–PVN regulation pathway that impacts both the upper GI prokinetic effects and anti-inflammatory effects of VNS. These findings identify the PVN as a central regulator of gastric motility and a key node for top-down control of the brain–vagal–gut axis, gating the effects of VNS.

## CRediT authorship contribution statement

**Chun-Huei Zou:** Investigation, Formal analysis. **Yu-Chieh Huang:** Investigation, Software, Formal analysis, Validation, Writing original draft & editing. **Wu-Lung Tseng:** Investigation, Formal analysis. **Ye-Ting Lin:** Investigation. **Ying-Jui Chao:** Conceptualization. **Yi-Jen Wu:** Conceptualization, Methodology, Resources, Supervision, Data Curation, Project administration, Writing – original draft, review & editing, Funding acquisition.

## Acknowledgements

We thank Prof. Kuei-Sen Hsu and Dr. Chin-Hao Chen, National Cheng Kung University, for their insightful comments and support of this study.

## Declaration of generative AI

During the preparation of this work, the authors used OpenAI to improve the language, readability, and clarity of the manuscript. After using this tool, the authors reviewed and edited the content as needed and take full responsibility for the content of the published article.

## Data Availability

The data supporting this manuscript is available upon request to the corresponding author.

---

**Supplementary Fig. 1.**
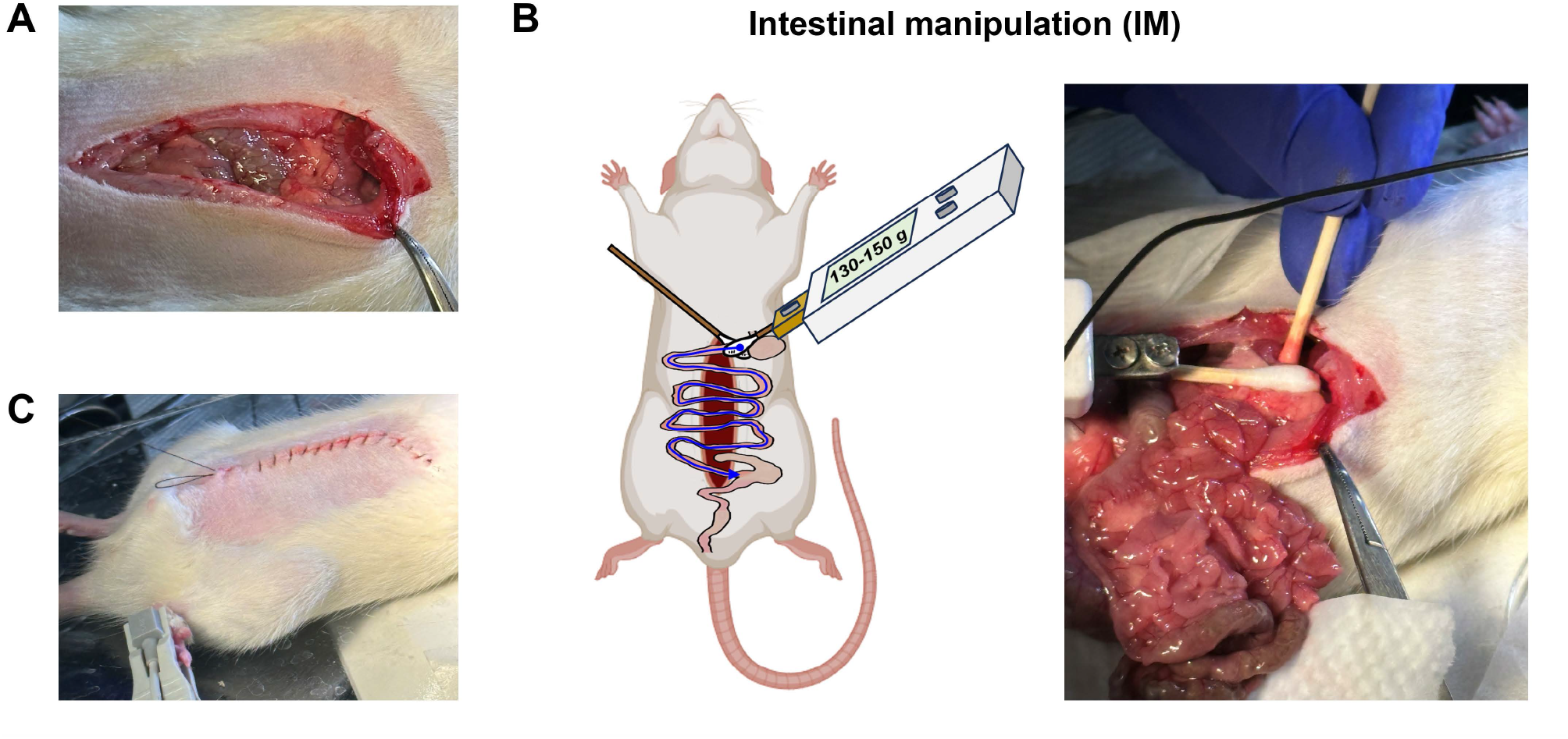
Intestinal manipulation (IM)-induced gastroparesis rats. **A,** Midline incision of the abdomen. **B,** Schematic of the IM procedure, in which the exteriorized small intestine was mechanically manipulated with saline-moistened cotton swabs at a standardized force of 130–150 g (left) and the intraoperative imaging (right). **C,** Closed abdominal incision line after IM.

**Supplementary Fig. 2.**
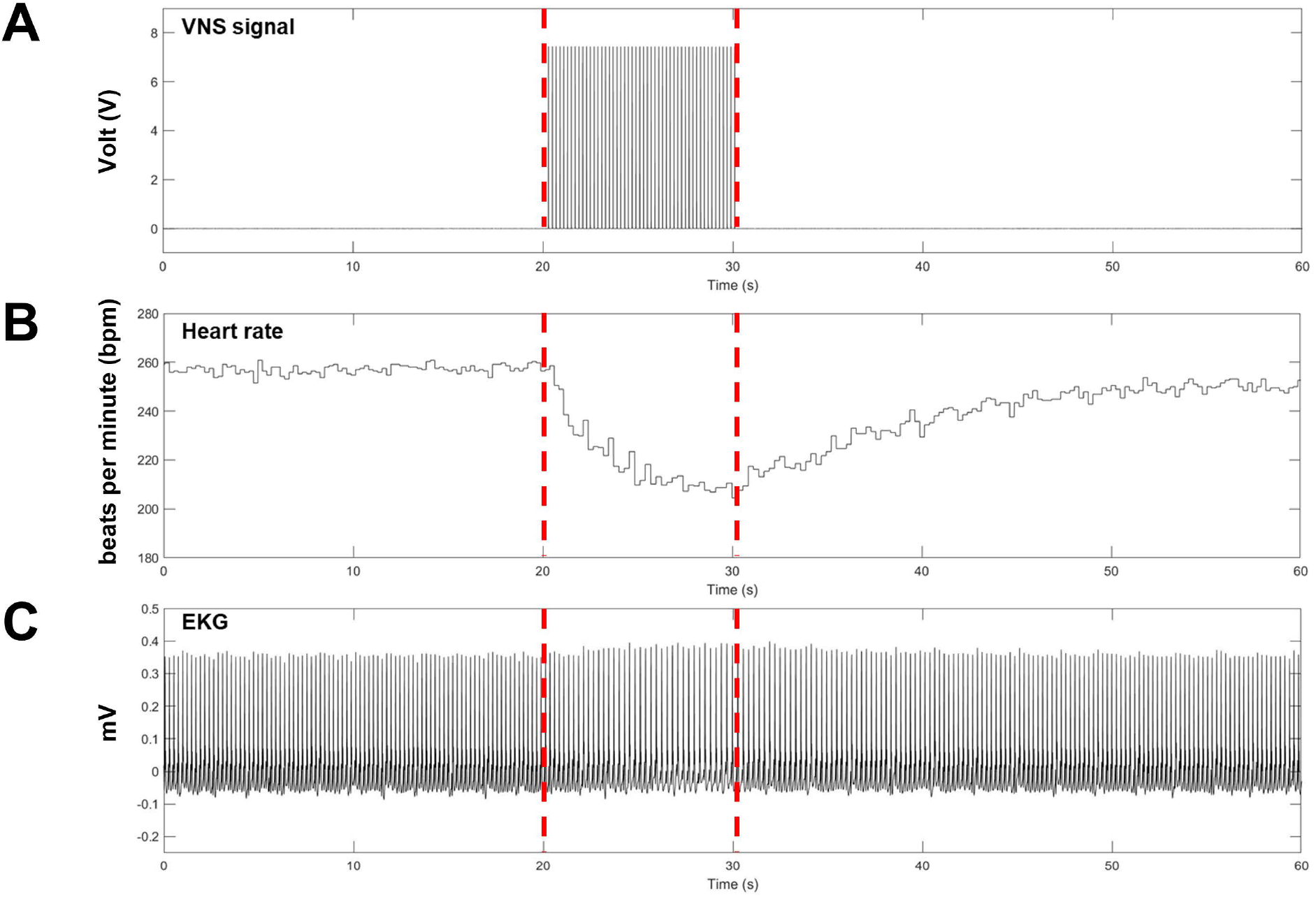
Heart rate changes during vagus nerve stimulation (VNS). Representative recording trace of electrocardiographic (EKG) before, during, and after a 10-s VNS train. **A**, VNS stimulation signal, **B,** heart rate, and **C,** EKG signals, revealing a decrease in heart rate during stimulation.

**Supplementary Fig. 3.**
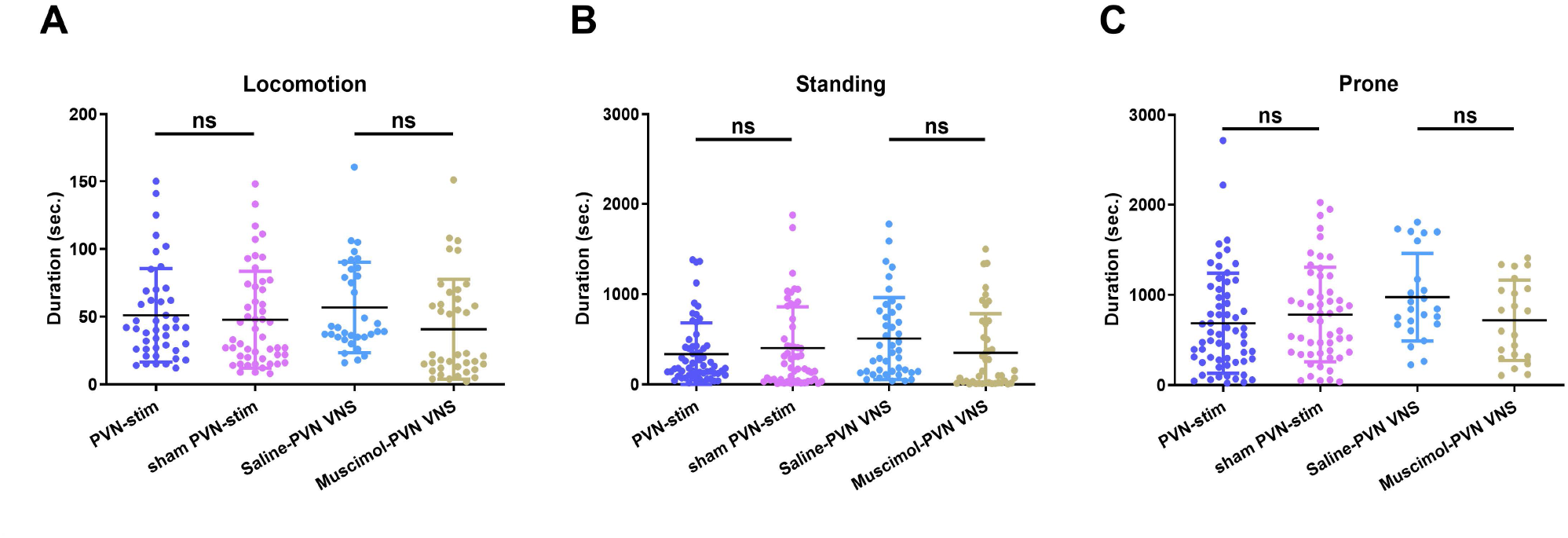
Post-surgical activity in gastroparesis rats following experimental treatments. Following PVN stimulation, sham stimulation, or VNS combined with intra-PVN injection of muscimol or saline, rats received oral gavage with 1.5 mL of FITC–dextran. Rats remained awake and were allowed to move freely in their home cages. Behavior was recorded for 3 h after gavage. **A**, Total duration of locomotion. **B,** Total duration of standing. **C,** Total duration of prone immobility. ns, non-significant.

## Supplementary Methods

### Supplementary Methods 1. FITC–dextran measurement of gastric emptying and gastrointestinal transit

Immediately after control laparotomy or intestinal manipulation for gastroparesis induction, rats received 1.5-mL of fluorescein isothiocyanate (FITC)-labeled dextran solution (FITC–dextran, Sigma-Aldrich, 46945-500MG-F, molecular weight 70,000, CAS 60842-46-8, 5 mg/mL) by oral gavage. After a 3-h transit period when animals were allowed to freely move in the home cage, they were deeply anesthetized with isoflurane (4%–5% for induction and 1%–3% for maintenance). The abdominal and thoracic cavities were opened, and 6 mL of whole blood was collected by cardiac puncture. The stomach was immediately ligated with 4-0 cotton suture at both the cardia and pylorus to prevent loss of luminal FITC–dextran that would confound gastric emptying quantification. The entire GI tract (stomach together with the small intestine, cecum and colon) was excised and temporarily stored under darkness while the animal was perfused transcardially with phosphate buffered saline (PBS) followed by 4% paraformaldehyde (PFA) in PBS to fix the brain for subsequent immunofluorescence analysis. The GI tract was then transferred to a fume hood, cleared of surrounding connective tissue, and straightened to measure its total length. Starting from the pylorus, the small intestine was divided into 10 equal-length segments (sb1–sb10) and the colon into three segments (col1–col3). Together with the stomach and cecum, the GI tract was thus divided into 15 sequential segments. The stomach was opened, and its luminal contents, together with those of each intestinal segment, were collected. Each segment was then flushed with 10 mL of Krebs–Henseleit buffer (KHB), and the luminal contents gently compressed with forceps into a collection tube along with the corresponding tissue segment. Samples were centrifuged at 1,500 × g for 5 min, and the supernatant was subjected to a second centrifugation at 11,000 × g for 10 min. The resulting supernatant was transferred to a 96-well plate, and fluorescence intensity measured using a microplate reader at excitation and emission wavelengths of 485 and 528 nm, respectively. Background fluorescence was subtracted using KHB blanks.

### Supplementary Methods 2. PVN recording and unit spike sorting Extracellular recording of PVN neuronal activity

Anesthetized rats were secured in a stereotaxic apparatus (RWD, CA, USA) and the skull exposed to perform a stereotactic surgery to target hypothalamic paraventricular nucleus (PVN). Eight custom-fabricated tungsten recording electrodes (99.95% purity, 50.8 μm bare diameter, HML-insulated, California Fine Wire Company, Grover Beach, CA, USA) were implanted bilaterally into the PVN (AP −1.8 mm, ML ±0.5 mm, DV −7.8 mm), four electrodes on the left side (channels 1–4) and four on the right side (channels 5–8). Electrodes were connected to a 16-channel connector (SenonLink Technology Co., Taiwan) and extracellular unit activities acquired using a multichannel recording system (Plexon, USA) with signal sampling at 20 kHz and band-pass filtering at 300–6000 Hz. Baseline neuronal activity was recorded before, during, and for several hours after VNS.

### PVN unit spike sorting

The extracellular unit spike recording signals from PVN during VNS or PVN stimulation were offline analyzed. For PVN stimulation, the stimulation artifacts were removed from the recording signals manually before sorting. Spike detection and initial clustering were performed using UltraMegaSort2000 (UMS2K, Hill et al., 2011) (https://neurophysics.ucsd.edu/software.php) and MATLAB. The spike detection threshold was defined as 2 times higher than the baseline amplitude, which was calculated from 2000 randomly selected data points per recording. After UMS2K sorting, the first three principal components (PC1, PC2, PC3) of each waveform were extracted to fit Gaussian mixture models (GMMs). K denotes the number of Gaussian components used to model the underlying data distribution, corresponding to the number of clusters tested for unit spikes in this study. For the recording PVN neuronal firing during VNS, four current intensities of VNS were applied (0.5, 1.0, 1.5, and 2.2 mA). At each intensity, there was one unit waveform sorted during the stimulation period. GMM models were therefore fitted with K = 1 to 4 to determine the number of statistically distinct subpopulations best described the units pooled across the four intensities. For direct stimulation recordings, units were recorded from both the left and right PVN and GMM models fit with K = 1 to 2 in order to determine whether left- and right-side units formed statistically distinct subpopulations. The Bayesian information criterion (BIC) was used for model selection, with lower values indicating a better-fitting model.

### Supplementary Methods 3. Serum collection and ELISA

Whole blood was collected by cardiac puncture as described in Supplementary Methods 1, placed on ice immediately without anticoagulant use, and centrifuged at 3,000×g for 15 min at 4°C. The supernatant was collected and centrifuged again at 10,000×g for 10 min for serum isolation. Serum samples were then aliquoted and stored at −80°C until analysis. Serum levels of interleukin-6 (IL-6), motilin, corticosterone, and ghrelin were measured in duplicate using commercially available ELISA kits (Supplementary Table 1) according to the manufacturers’ instructions. Concentrations were calculated from standard curves generated for each assay.

### Brain section preparation and immunofluorescence

Animals were deeply anesthetized and transcardially perfused with normal saline, followed by 4% PFA. Brains were removed, post-fixed in 4% PFA overnight at 4°C, and cryoprotected in 30% sucrose solution until the tissue sank. Coronal brain sections were cut into 40-μm thick sections on a sliding microtome (SM2010R, Leica Biosystems) over dry ice, then washed in wash buffer (0.1% Triton X-100 in 1× PBS) and blocked in antibody diluent with background reducing components (S3022, Dako, Agilent). Sections were incubated overnight at 4°C with primary antibodies against c-Fos (1:1000, mouse monoclonal, Abcam ab208942) and either NeuN (1:500, rabbit monoclonal, Abcam ab177487) or CRH (1:1000, rabbit polyclonal, Abcam ab8901). Sections were washed in wash buffer and incubated for 2 h at room temperature with Alexa Fluor 488 goat anti-mouse Ig (1:1000, A-11001, Thermo Fisher) for c-Fos, and Alexa Fluor 568 goat anti-rabbit Ig (1:1000, A-11011, Thermo Fisher) for NeuN or CRH, then mounted using DAPI-containing mounting medium (ab104139, Abcam). Images were acquired using a confocal laser scanning microscope (FV3000, Olympus). NeuN-positive and NeuN/c-Fos double-positive cells (Fig. 2), as well as CRH-positive, c-Fos-positive, and CRH/c-Fos double-positive cells (Fig. 7E) were quantified within anatomically defined regions by investigators blinded to treatment groups, using a panoramic tissue cytometry system (TissueFAXS, TissueGnostics) and TissueQuest software. For each animal, cell counts were averaged across 1 to 3 sections to yield a single value per animal.

### Gastrointestinal motilin receptor (MTLR) and myeloperoxidase (MPO) immunohistochemistry

Gastrointestinal tissue samples were fixed in 10% neutral buffered formalin, paraffin-embedded, sectioned at 4 μm, deparaffinized, rehydrated, and heated to 95–98°C in Dako Target Retrieval Solution (citrate buffer, pH 6.0) for 1 h using a heated stirrer. Endogenous peroxidase activity was blocked with 3% H₂O₂ in ddH₂O for 15 min, followed by incubation with Dako blocking buffer (S3022) for 1 h at room temperature. For MTLR immunohistochemistry, sections were incubated with an anti-motilin receptor antibody (MyBioSource, #MBS2522198, 1:100) overnight at 4°C, washed, and incubated with Dako anti-rabbit secondary antibody (K4003) for 1 h at room temperature. Immunolabeling was visualized using 3,3′-diaminobenzidine (DAB). For MPO immunohistochemistry, sections were incubated with an anti-MPO antibody (rabbit polyclonal, ab9535, Abcam, 1:100) overnight at 4°C, washed, and developed using the PolyDetector Liquid DAB HRP Brown kit (BSB-0016, Bio SB). All sections were counterstained with hematoxylin, dehydrated, cleared, and mounted. Brightfield images were acquired using an inverted microscope (IX73, Olympus) and analyzed using ImageJ (Fiji). Images were converted to grayscale, and a fixed threshold was applied to identify MTLR- or MPO-positive cells. Positive cells within regions of interest were counted and normalized to tissue area in per mm².

---

**Supplementary Table 1.**

| **Reagent/assay** | **Manufacturer** | **Catalog no.** | **Dilution/concentration** |
| --- | --- | --- | --- |
| FITC–dextran | Sigma-Aldrich | 46945-500MG-F | 5 mg/mL |
| Muscimol | Sigma-Aldrich, USA | M1523 | 0.5 nmol/70 nL |
| Blocking buffer / antibody diluent | Dako (Agilent) | S3022 | N/A |
| NeuN antibody | Abcam | ab177487 | 1:500 |
| CRH antibody | Abcam | ab8901 | 1:1000 |
| c-Fos antibody | Abcam | ab208942 | 1:1000 |
| Alexa Fluor 568 goat anti-rabbit secondary antibody | Thermo Fisher | A-11011 | 1:1000 |
| Alexa Fluor 488 goat anti-mouse secondary antibody | Thermo Fisher | A-11001 | 1:1000 |
| DAPI mounting medium | Abcam | ab104139 | N/A (ready-to-use) |
| Motilin receptor antibody | MyBioSource | MBS2522198 | 1:100 |
| Anti-rabbit secondary antibody / DAB detection (MTLR) | Dako (Agilent) | K4003 | N/A (ready-to-use) |
| MPO antibody | Abcam | ab9535 | 1:100 |
| PolyDetector Liquid DAB HRP Brown kit (MPO detection) | Bio SB | BSB-0016 | N/A (ready-to-use) |
| IL-6 ELISA | BioLegend (LEGEND MAX™) | 437107 | - |
| Motilin ELISA | MyBioSource | MBS2099150 | - |
| Ghrelin ELISA | Elabsience | E-EL-R0842 | - |
| Corticosterone ELISA | Cayman Chemical | 501320 | - |

Details of the materials used, including the manufacturer, catalog number, and dilution or concentration of the reagent or assay.

**Supplementary Table 2.**

| Fig. | Label | Unit Spike Feature (Mean ± SD) | | | GMM | | | |
| --- | --- | --- | --- | --- | --- | --- | --- | --- |
| | | Amplitude<br>($\mu$ V) | Half Width<br>(ms) | Firing Rate<br>(Hz) | BIC<br>(K = 1) | BIC<br>(K = 2) | BIC<br>(K = 3) | BIC<br>(K = 4) |
| Fig. 2H and 2I | 0.5 mA VNS | 43.18 ± 11.60 | 0.147 ± 0.050 | 14.80 ± 0.92 | -26431.6 | -26367.8 | -26302.7 | -26230.6 |
| | 1.0 mA VNS | 42.42 ± 10.82 | 0.151 ± 0.057 | 16.29 ± 1.01 | | | | |
| | 1.5 mA VNS | 42.93 ± 11.05 | 0.147 ± 0.056 | 17.52 ± 1.06 | | | | |
| | 2.2 mA VNS | 43.22 ± 10.72 | 0.154 ± 0.066 | 20.31 ± 1.07 | | | | |
| Fig. 3B and 3C | PVN-stim, rec. left PVN | 21.31 ± 4.12 | 0.351 ± 0.090 | 20.27 ± 1.92 | -8436.1 | -8128.4 | - | |
| | PVN-stim, rec. right PVN | 21.79 ± 4.45 | 0.343 ± 0.083 | 18.78 ± 1.96 | | | | |

**Supplementary Table 3.**

| **Fig. 2J. VNS condition** | **Firing rate (Mean ± SD)** | **P value** |
| --- | --- | --- |
| Pre-stim vs. 0.5 mA (on) | 3.31 ± 1.15 vs. 14.80 ± 0.92 | <0.0001 |
| Pre-stim vs. 1.0 mA (on) | 3.31 ± 1.15 vs. 16.29 ± 1.01 | <0.0001 |
| Pre-stim vs. 1.5 mA (on) | 3.31 ± 1.15 vs. 17.52 ± 1.06 | <0.0001 |
| Pre-stim vs. 2.2 mA (on) | 3.31 ± 1.15 vs. 20.31 ± 1.07 | <0.0001 |
| Post-stim (1) vs. 0.5 mA (on) | 3.28 ± 0.67 vs. 14.80 ± 0.92 | <0.0001 |
| Post-stim (1) vs. 1.0 mA (on) | 3.28 ± 0.67 vs. 16.29 ± 1.01 | <0.0001 |
| Post-stim (1) vs. 1.5 mA (on) | 3.28 ± 0.67 vs. 17.52 ± 1.06 | <0.0001 |
| Post-stim (1) vs. 2.2 mA (on) | 3.28 ± 0.67 vs. 20.31 ± 1.07 | <0.0001 |
| Post-stim (2) vs. 0.5 mA (on) | 2.56 ± 0.30 vs. 14.80 ± 0.92 | <0.0001 |
| Post-stim (2) vs. 1.0 mA (on) | 2.56 ± 0.30 vs. 16.29 ± 1.01 | <0.0001 |
| Post-stim (2) vs. 1.5 mA (on) | 2.56 ± 0.30 vs. 17.52 ± 1.06 | <0.0001 |
| Post-stim (2) vs. 2.2 mA (on) | 2.56 ± 0.30 vs. 20.31 ± 1.07 | <0.0001 |
| 0.5 mA (on) vs. 1.0 mA (on) | 14.80 ± 0.92 vs. 16.29 ± 1.01 | 0.0134 |
| 1.0 mA (on) vs. 1.0 mA (off) | 16.29 ± 1.01 vs. 13.60 ± 0.33 | <0.0001 |
| 1.5 mA (on) vs. 1.5 mA (off) | 17.52 ± 1.06 vs. 13.28 ± 0.70 | <0.0001 |
| 0.5 mA (on) vs. 1.5 mA (on) | 14.80 ± 0.92 vs. 17.52 ± 1.06 | <0.0001 |
| 1.5 mA (on) vs. 2.2 mA (on) | 17.52 ± 1.06 vs. 20.31 ± 1.07 | <0.0001 |
| 2.2 mA (on) vs. 2.2 mA (off) | 20.31 ± 1.07 vs. 13.00 ± 0.71 | <0.0001 |
| 0.5 mA (on) vs. 2.2 mA (on) | 14.80 ± 0.92 vs. 20.31 ± 1.07 | <0.0001 |
| 1.0 mA (on) vs. 2.2 mA (on) | 16.29 ± 1.01 vs. 20.31 ± 1.07 | <0.0001 |

